# FunCoup 6: advancing functional association networks across species with directed links and improved user experience

**DOI:** 10.1101/2024.09.13.612391

**Authors:** Davide Buzzao, Emma Persson, Dimitri Guala, Erik L.L. Sonnhammer

## Abstract

FunCoup 6 (https://funcoup6.scilifelab.se/, will be https://funcoup.org after publication) represents a significant advancement in global functional association networks, aiming to provide researchers with a comprehensive view of the functional coupling interactome. This update introduces novel methodologies and integrated tools for improved network inference and analysis. Major new developments in FunCoup 6 include vastly expanding the coverage of gene regulatory links, a new framework for bin-free Bayesian training, and a new website. FunCoup 6 integrates a new tool for disease and drug target module identification using the TOPAS algorithm. To expand the utility of the resource for biomedical research, it incorporates pathway enrichment analysis using the ANUBIX and EASE algorithms. The unique comparative interactomics analysis in FunCoup provides insights of network conservation, now allowing users to align orthologs only or query each species network independently. Bin-free training was applied to 23 primary species, and in addition networks were generated for all remaining 618 species in InParanoiDB 9. Accompanying these advancements, FunCoup 6 features a new redesigned website, together with updated API functionalities, and represents a pivotal step forward in functional genomics research, offering unique capabilities for exploring the complex landscape of protein interactions.

**GRAPHICAL ABSTRACT:** 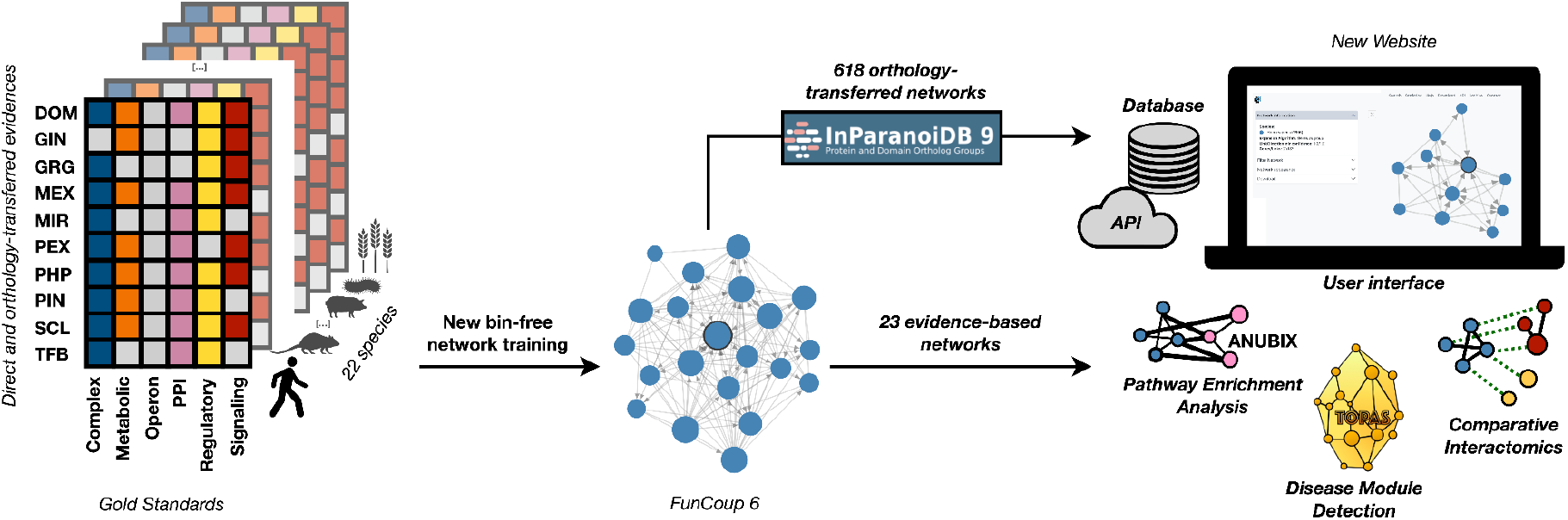

## INTRODUCTION

Elucidating protein interactions is fundamental for unraveling complex cellular processes. Current methodologies, ranging from precise but costly low-coverage experiments to cost-efficient yet error-prone high-throughput techniques like yeast two-hybrid, yield diverse interaction types, from direct physical to regulatory interactions. Existing databases like BIND (Salwinski et al. 2004), BioGRID (Oughtred et al. 2021), DIP (Salwinski et al. 2004), IntAct (Del Toro et al. 2022), ENCODE (ENCODE Project Consortium 2012; Luo et al. 2020), MINT (Licata et al. 2012), SIGNOR (Lo Surdo et al. 2023), Reactome (Gillespie et al. 2022), HPRD (Keshava Prasad et al. 2009), PhosphoSitePlus (Hornbeck et al. 2015), and HINT (Das and Yu 2012) store primary interaction data but may lack the comprehensive integration needed for a holistic understanding of the interactome. A robust framework for data integration is crucial for minimizing errors and capturing a comprehensive view of cellular processes and connecting them to functions and phenotypes.

Several frameworks, including FunCoup (Persson et al. 2021), STRING (Szklarczyk et al. 2023), HumanNet (Kim et al. 2022), PrePPI (Petrey et al. 2023), GeneMania (Warde-Farley et al. 2010), HumanBase (Greene et al. 2015), IMP (Wong et al. 2015), I2D (Kotlyar et al. 2016), and ConsensuspathDB (Herwig et al. 2016), employ different machine learning techniques to integrate multi-omics data and represent interactomes with genes or proteins as nodes linked by functional associations. Researchers leverage the versatile capabilities of functional association networks across diverse domains, including hypothesis generation for both small-scale and large-scale data analyses, and contributing to the advancement of third party resources. The FunCoup database is built with a unique redundancy-weighted naïve Bayesian approach, combining 10 diverse data types and incorporating orthology transfer, to infer functional associations for 23 species. FunCoup has demonstrated superior performance in identifying disease genes using ORPHANET (Weinreich et al. 2008) gene sets (Persson et al. 2021) and ranked among the top three best performers in a benchmark of 46 networks (Wright et al. 2024). It facilitates hypothesis generation by revealing intricate gene relationships, as seen e.g. in studies exploring the roles of genes post-Malat1 knockdown (Aslanzadeh et al. 2024), uncovering co-expression patterns in disease contexts like CHARGE syndrome (Carrara et al. 2024), and identifying relevant clusters in non-ossifying fibroma (Souchelnytskyi 2023). FunCoup was integrated into a deep learning framework for predicting drug-target interactions (Wang et al. 2023), and in the development of the DiffBrainNet resource (Gerstner et al. 2022). FunCoup has further facilitated pathway enrichment analysis through tools like PathBIX, e.g. to investigate TP53 signaling disruptions in osteosarcoma (Saba et al. 2024) and the exploration of ALS-related gene networks (Mei et al. 2024).

As the underlying data evolves, functional association networks require continuous framework updates. Previous versions of FunCoup, as well as other functional association networks, suffer from arduous training methodologies and coarse resolution due to binning techniques, resulting in low accuracy in link prediction and poor reproducibility. Furthermore, existing functional association databases generally lack gene regulatory interactions and are restricted in terms of species representation, creating gaps in the understanding of complex biological processes. Many resources have user interface limitations that impede the effective integration and use of advanced analytical tools. FunCoup 6 addresses these limitations by incorporating new methodologies and tools to enhance network inference and analysis. Key advancements include significantly expanded gene regulatory link coverage, a novel bin-free naïve Bayesian training framework, and a redesigned website. For 13 species, the number of regulatory links has increased substantially, with the human network alone featuring around half a million directed links, deepening our understanding of transcriptional regulation.

In addition to the MaxLink tool (Guala, Sjölund, and Sonnhammer 2014) for candidate gene prioritization, FunCoup 6 now includes the TOPAS algorithm (Buzzao et al. 2022) for identifying disease and drug target network modules. The resource is further enhanced with integrated KEGG (Kanehisa et al. 2014) pathway enrichment analysis utilizing the ANUBIX (Castresana-Aguirre and Sonnhammer 2020) and EASE (Hosack et al. 2003) algorithms, enhancing its application in biomedical research. The unique comparative interactomics feature in FunCoup, which provides insights into network conservation across species, has been enhanced with a new search mode. This mode performs a network expansion in the other species with orthologs to the query, to identify interactors independently in each species. The new bin-free training was applied to 23 primary species, and networks for an additional 618 species in InParanoiDB 9 (Persson and Sonnhammer 2023) were generated using a new orthology-based method. These improvements are complemented by a redesigned website and extended API functionalities, enhancing user accessibility and experience.

## RESULTS

### Bin-free network training and updated data

In order to derive functional association likelihoods from high-throughput data and gold standards, FunCoup 6 employs a new bin-free naïve Bayesian framework that uses kernel density estimation and polynomial regression to compute log-likelihood ratios, and a new weighted redundancy schema to extract a Final Bayesian Score (FBS) for each link and gold standard (Supplementary Figure 3). This method requires the data size and regression quality to meet specific thresholds for reliable results. A total of 2969 combinations of gold standards and evidences met these requirements, accounting for 34% of all possible combinations across all species (Supplementary Figure 4). The majority of unsuccessful training was due to lack of data (i.e. too few gold standard examples to extract likelihoods).

In FunCoup 6, we renamed the evidence QMS to PEX and updated the data sources for the evidences PEX, PHP, PIN, and SCL (Supplementary Table 1). To achieve broader coverage and more representative expression data for healthy individuals, we utilized different databases for GIN, MEX, MIR and TFB, and integrated a new evidence GRG for regulatory interactions. Due to the new training framework requiring continuous data, we made several changes to the evidence scoring methods, which are summarised in Supplementary Table 1. Further details can be found in the Supplementary Materials.

The new confidence estimation method in FunCoup calculates Positive Predictive Value (PPV) across varying FBS by using gold standard links as positive examples and an equal number of random pairs as negative examples, then fitting a logistic function to the PPV vs FBS relation to provide an empirical measure of the confidence of functional associations. This method requires data and regression to meet specific criteria for accuracy (Supplementary Figure 5). A total of 57 gold standards have met these requirements, accounting for 83% of all networks across all species (Supplementary Figure 6). The most common reason for failed PPV extraction was poor fitting (*i*.*e*. logistic regression fit with *R*^2^ < 0. 9).

### Evidence-based networks for 23 species

In FunCoup 5 we provided functional association networks for 22 species, composed of 17 eukaryotes, 2 bacteria, 2 archaea and the virus SARS-CoV-2. In FunCoup 6 we added one more bacterial species, *Mycobacterium tuberculosis*, reaching a total of 23 species. SARS-CoV-2 was built as a host-virus interactome together with human proteins, and was updated to the latest UniProt reference proteome and IrefIndex release of interactions, resulting in a total of 2,879 links, of which 2,833 are links between 17 reference proteins and 1,924 human proteins, and 46 are links between viral proteins only.

The proteome coverage of the networks for each species can be seen in Figure 1A. The coverage has increased from FunCoup 5 for all species except *C. intestinalis* which decreased by thirteen percentage points. The proteome coverage exceeds 50% for all species except *S. solfataricus* and *M. tuberculosis* where it remains relatively low (37-44%), due to insufficient data for these less studied species.

**Figure 1.**
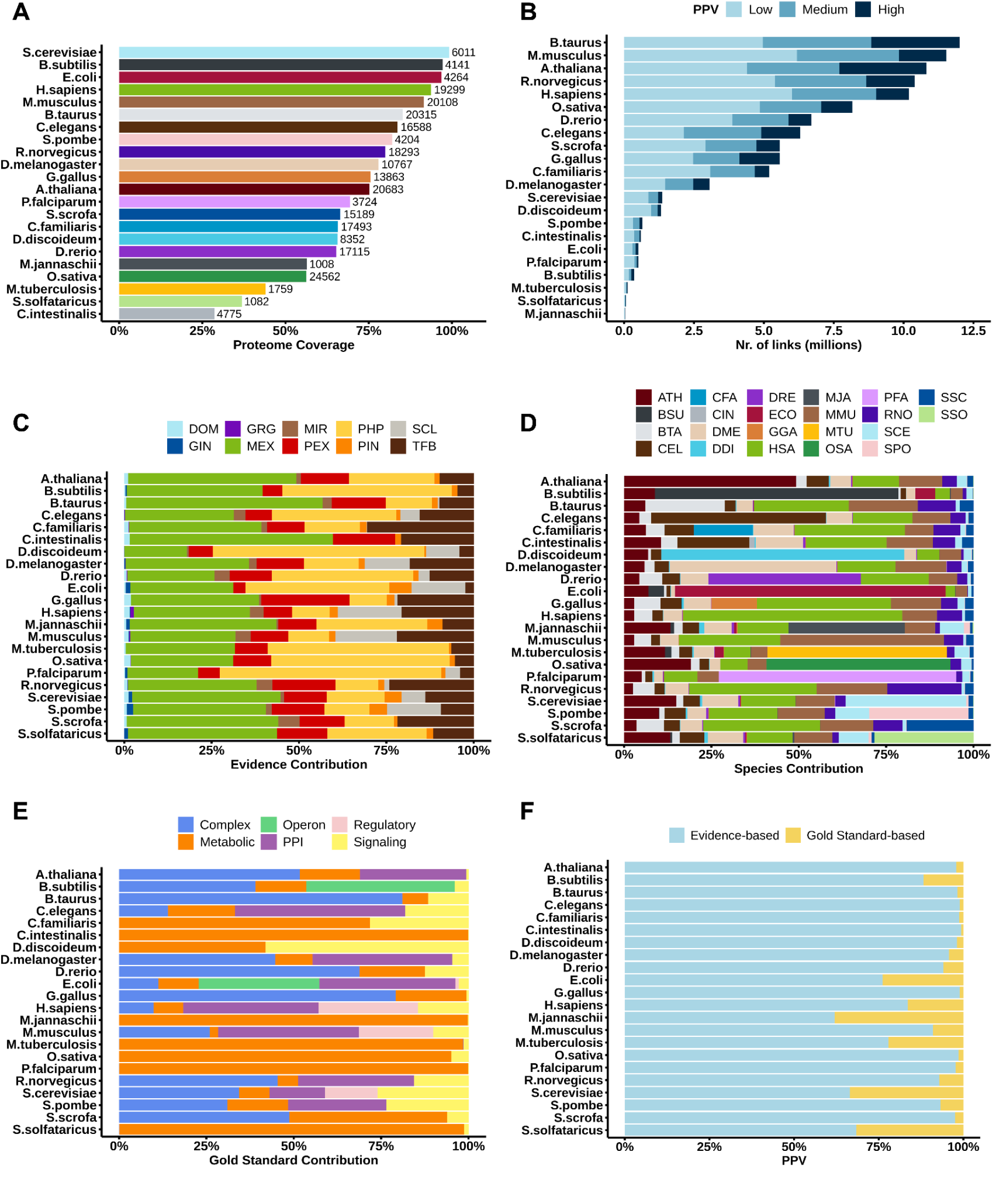
FunCoup 6 statistics. (A) The proteome coverage of the networks for each of the species in FunCoup 6. (B) The number of links for different PPV confidence cutoffs for each of the species in FunCoup 6. Low PPV confidence is PPV between 0.85 and 0.9, medium is PPV between 0.9 and 0.95, high is PPV above 0.95. (C) Contribution of evidence types to the networks of each FunCoup 6 species. The sum of LLRs is normalized so it sums to 1 per species. Evidence data types are: DOM: domain interactions, GIN: genetic interaction profile similarity; GRG: gene regulation; MEX: mRNA co-expression; MIR: co-miRNA regulation by shared miRNA targeting; PEX: protein co-expression; PHP: phylogenetic profile similarity; PIN: protein interaction networks; SCL: sub-cellular co-localization; TFB: shared transcription factor binding. (D) Evidence contribution from other species in each FunCoup 6 network. (E) Contribution of Gold Standard to the networks of each FunCoup 6 species. The number of links is normalized so it sums to 1 per species. (F) The percentage of links with Gold Standard- and Evidence-based PPV.

The total number of functional associations has increased from 60,999,771 in FunCoup 5, to 101,054,256 in FunCoup 6. The distribution of links over different confidence score cutoffs for each species can be found in Figure 1B. For a majority of the species, the number of links in the network has increased in comparison to FunCoup 5, although for 3 species (*C. familiaris, C. intestinalis, and D. rerio*) we can see a decrease in the number of links.

The positive relative contribution of evidence types to the total FBS per species can be seen in Figure 1C. While the overall distribution of evidence types remains similar, MEX has decreased in its contribution compared to previous releases due to more stringent inclusion criteria, with PHP now taking a more prominent role due to its large coverage. PIN had limited coverage and a highly discretized score distribution in FunCoup 5, resulting in predominantly low or high scores, providing only a small contribution in all species except in *E. coli, S. cerevisiae, and S. pombe*. In FunCoup 6, the PIN contribution remains small but is more uniformly distributed across species, with much smaller contributions in those three species. TFB has been notably boosted compared to the previous release. Similarly, GIN and DOM have also received enhancements, while MIR remains largely unchanged. However, these three evidences play a minimal contributing role overall.

Figure 1D illustrates the positive relative contribution of orthology transfer from other species to the total FBS of each network. Similar to previous FunCoup versions, the majority of species obtain more than 50% of their evidence through orthology transfer from other species.

For each species, the final FunCoup network corresponds to the union of the gold standard networks, keeping the maximum PPV. Only links with PPV ≥ 0.85 are kept. Figure 1E demonstrates that the contributions of various gold standards, such as Complex, Metabolic, PPI, and Signaling, vary across different species. While some species rely heavily on one or two types of gold standards, others display a more balanced mix. Operon was only used for bacteria. A new Regulatory gold standard was added for *H. sapiens, M. musculus, E. coli* and *S. cerevisiae*.

Out of the over 100 million links, 7,383,393 are Gold Standard links. The updated FunCoup scoring system integrates Gold Standard-based PPV by assigning values based on the number of supporting Gold Standards, prioritizing manually curated evidence over lower Evidence-based PPV values for higher prediction accuracy. Figure 1F shows the percentage of links with Gold Standard- and Evidence-based PPV. The majority of links (65-99%) were assigned an Evidence-based PPV.

### Directed links

In FunCoup 5 we introduced directed regulatory links at a very modest scale in human, inferred from ChIP-seq data, to denote transcription factor to target gene binding. In FunCoup 6, significant advancements were made in expanding regulatory networks across 13 species, for a sum of 964,037 directed links with positive GRG (Figure 2A). Positive GRG is sufficient to make the link directed, and it can be found for any gold standard. In the human network, there are 838 transcription factors interacting with 14,989 target genes, for a total of 501,720 directed links, 450 times more than in FunCoup 5. As an example, Figure 2B shows a subnetwork around the transcription factor JUN, known to play a significant role in cellular stress responses, apoptosis, and proliferation (Behrens, Sibilia, and Wagner 1999). Among its highest confidence interactions, links to MAPK8, ESR1, CREB5 and CTNNB1 were previously reported in TRRUST and/or RegNetwork. Other interactions, like those with ATF2, MECOM and SPI1, are highly supported by the GRG evidence, suggesting further complexity in the JUN regulatory network, for which support in the literature exists (Liu et al. 2006, Shen et al. 2016). Overall, the links in the network were inferred with Regulatory and PPI as the strongest gold standard and PIN as the strongest evidence type.

**Figure 2.**
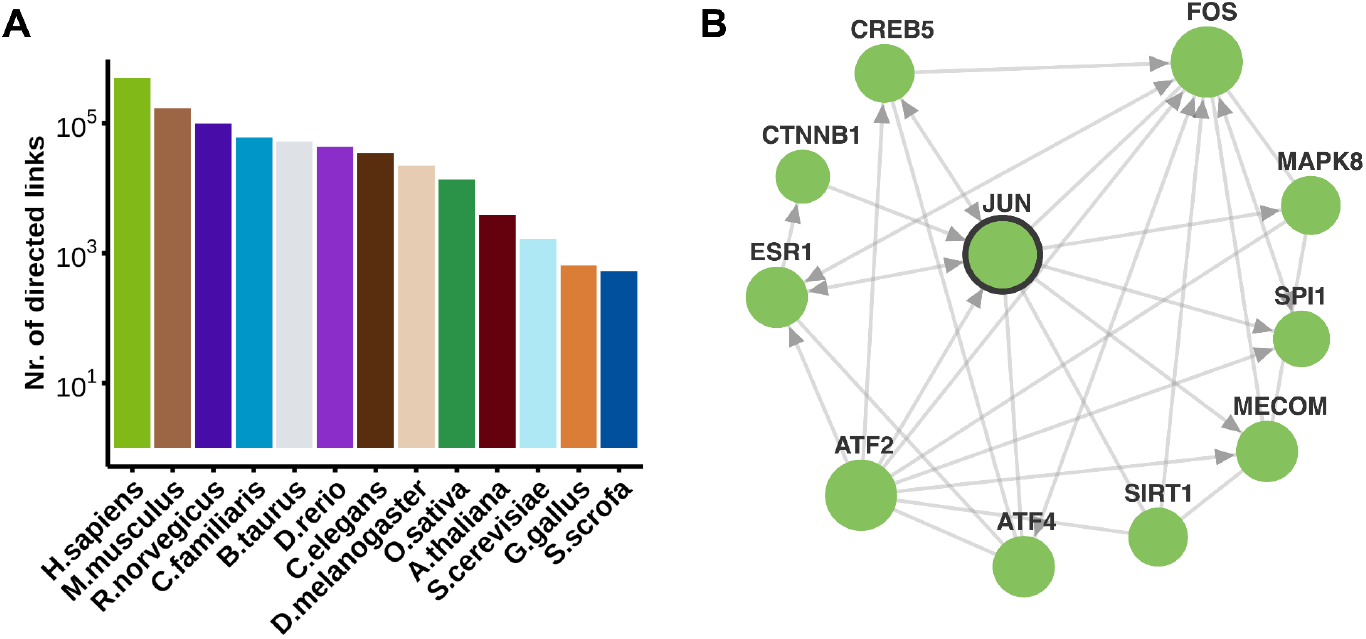
Directed links in FunCoup 6 and a JUN regulatory subnetwork in human. (A) The number of directed links in 13 FunCoup species. (B) A *JUN* FunCoup subnetwork, with 11 nodes and 30 links, of which 23 are directed to exhibit gene regulation. The database query (link) was performed on the website with the following parameters: query gene “JUN”; links and direction confidence threshold of 0.95 and 1; expansion depth 1; max 10 neighbors. Genes were searched as a group, with the prioritization of neighbors option activated. In the network illustration: the query gene is highlighted with a bold outline, the size of the nodes is proportional to the degree of connectivity. and nodes are colored in green using a new website feature.

### Network search algorithms

There are four algorithms in FunCoup 6 to search the network to expand a set of query genes to neighboring genes: Independent, Group, MaxLink, and TOPAS. To exemplify these, we applied them to a query of *Biliary liver cirrhosis* genes, as described in a published collection of 70 disease gene sets (Ghiassian, Menche, and Barabási 2015). This gene set (21 genes) is highly disconnected in the human network (6 links, Supplementary Figure 7A) when we use high-confidence links in FunCoup (PPV ≥ 0.95), indicating that many relevant genes might be missing from the query.

In the independent search (Supplementary Figure 7B), each gene is queried separately, expanding the network by adding the top most confident interactors for each gene. This method generally results in a rapid increase in network size. The group search method (Supplementary Figure 7C), especially when the “common neighbor prioritization” option is activated, tends to expand the network to a lesser degree. It adds top interactors with the highest number of interactions above the cutoff to the entire query group. This approach identified immediate neighbors that interact with multiple query genes, thus forming a cohesive subnetwork, but not as homogeneously throughout the genes.

The MaxLink algorithm (Supplementary Figure 7D) prioritizes genes based on the statistical significance of the connections to the query. However, it here struggled to identify new candidates because the input genes are largely disconnected and have few shared links to other genes. In contrast, the TOPAS algorithm (Supplementary Figure 7E) excelled in this context. TOPAS identified 11 connectors for 20 seed/query genes, 2 of which (*i*.*e*. B2M and PDGFR) were previously shown to be relevant in liver cirrhosis (Buzzao et al. 2022), demonstrating its utility in constructing biologically connected disease modules from a sparse and incomplete gene set.

#### Comparative interactomics analysis

One of the unique functionalities of FunCoup is its comparative interactomics search to investigate network conservation between species. The redesigned comparative interactomics allows users to either align orthologs of the expanded query network only, or to search each species with orthologs to the query and expand separately in each species. By default, only species-specific evidence is used to avoid conservation from orthology-based evidence. In Figure 3 we illustrate comparative interactomics by searching in *C. elegans* for the network conservation of Presenilin-1 (PSEN1), Presenilin-2 (PSEN2), and Filamin B (FLNB), three genes implicated in Alzheimer’s disease (AD) in human (Duff et al. 1996). PSEN1 and PSEN2 are involved in the processing of amyloid precursor protein and FLNB in cytoskeletal organization. PSEN1 and PSEN2 are orthologs to sel-12, and FLNB to fln-1. Of particular interest are “rectangular” constellations, which consist of two functional associations and two ortholog links, here formed by PSEN1-CRK and sel-12-ced-2. CRK has not previously been associated with AD, but it is a signaling protein involved in cell adhesion which is associated to the physiological functions of amyloid precursor proteins and the formation of neurotoxic amyloid-ß peptide aggregates, which are considered to play a central role in the etiology of the disease (Pfundstein, Nikonenko, and Sytnyk 2022; Leshchyns’ka and Sytnyk 2016; Wennström and Nielsen 2012). The ortholog in *C. elegans*, ced-2, is involved in cell death which is a hallmark of AD. Thus, investigating ced-2 in the worm offers advantages such as using its genetic tools for precise studies, gaining insights into the role of CRK in AD pathways involving PSEN1, and revealing evolutionarily conserved mechanisms relevant to disease research.

**Figure 3.**
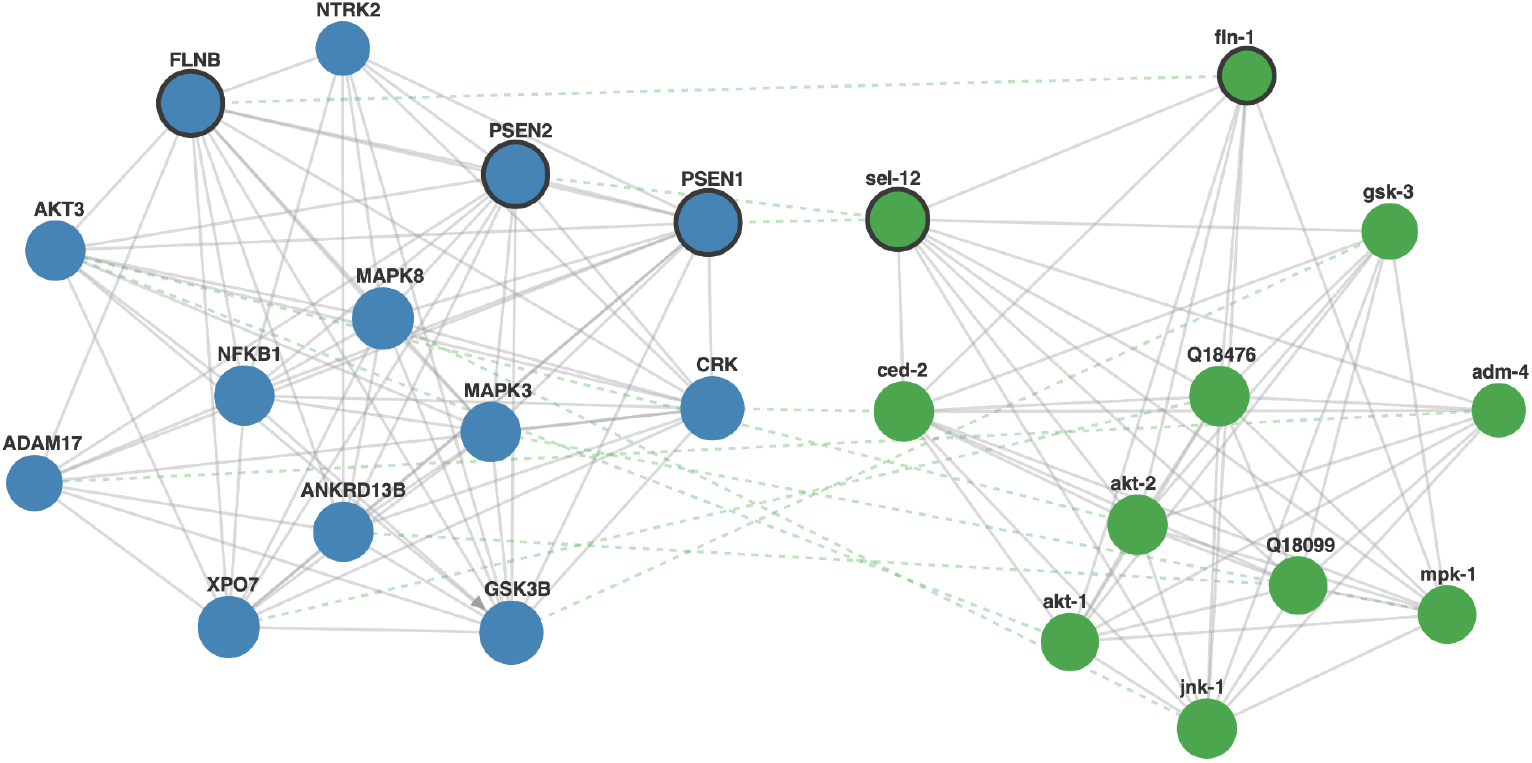
Comparative interactomics analysis of an Alzheimer’s disease related gene set. Mixed subnetworks of the genes PSEN1, PSEN2, and FLNB in *H. sapiens* (blue) and *C. elegans* (green) showing functional (gray solid lines) and ortholog links (green dashed lines). The database query (link) was performed via the website with the following parameters: query genes “PSEN1,PSEN2,FLNB”; query species *H. sapiens* and comparative species *C. elegans*; links and direction confidence threshold of 0.5 and 1; expansion depth 1; max 10 neighbors. Genes were searched as a group, with the prioritization of neighbors option activated. In the network illustration: query genes are highlighted with bold outlines, and the size of the nodes is proportional to the degree of connectivity.

### Network benchmark

To compare the performance of FunCoup 6 with FunCoup 5, HumanNet v3, and STRING v12, we split disease-associated genes from 110 distinct ORPHANET gene sets into two halves and assessed their recovery using Random Walk with Restart (RWR), measuring Performance Gain (PG) across five scenarios with varying numbers of high-confidence links (Figure 4A). FunCoup 6 shows superior performance compared to the other networks. Despite a larger standard deviation, FunCoup 6 exhibits higher median performance gain. The full distribution of PG at the various cutoffs is shown in Supplementary Figure 8.

**Figure 4.**
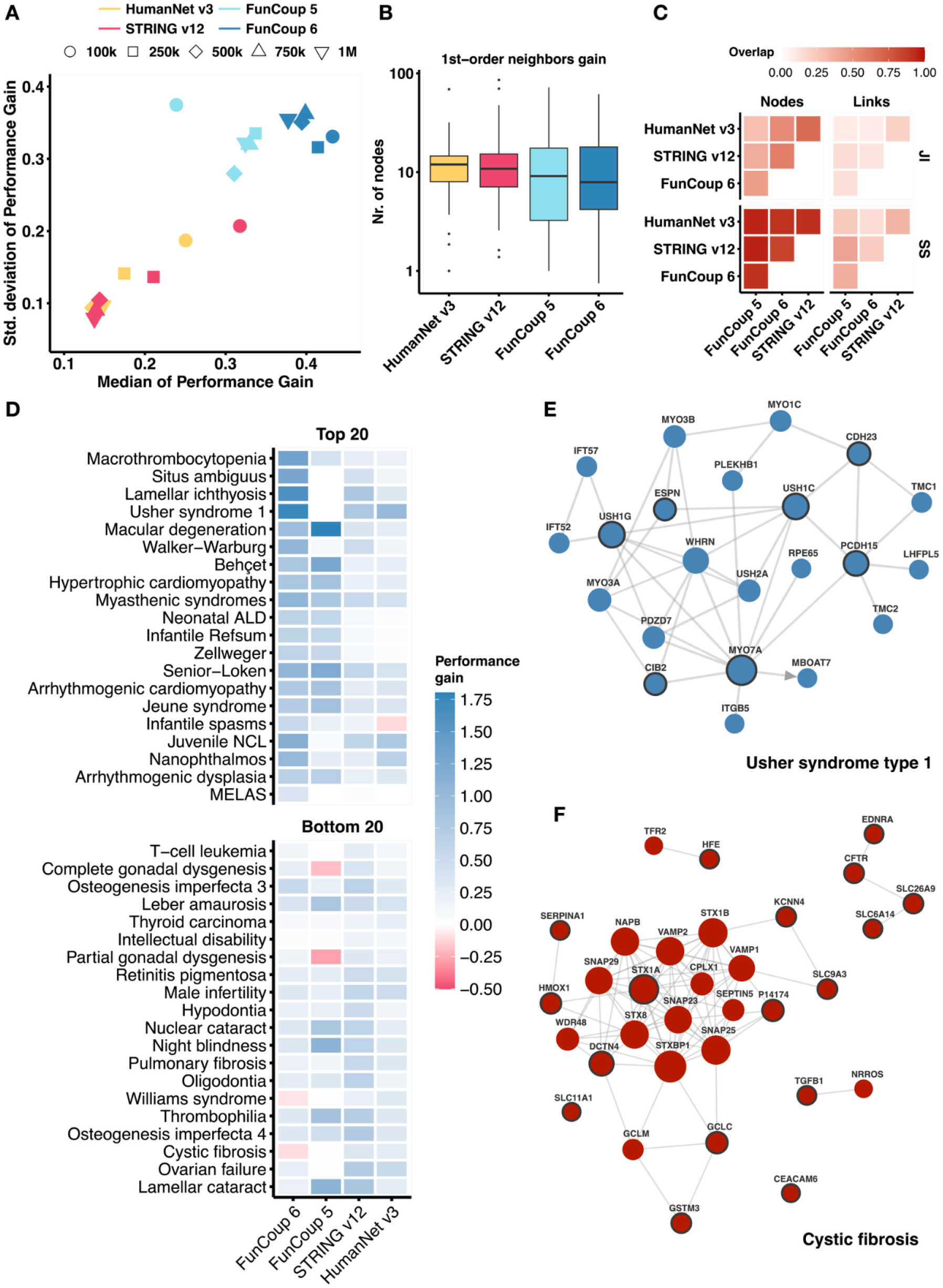
Performance evaluation of disease gene sets from ORPHANET across varying network configurations. (A) Performance gain is quantified as the difference in Area Under the Receiver Operating Characteristic curve (AUROC) between real networks and their null counterparts. The assessment was done using a top fixed number of links: 100,000, 250,000, 500,000, 750,000, and 1.000.000, and summarized in a scatter plot with colors and shapes indicating networks and links cutoff. (B) The distributions of ORPHANET gene set sizes after mapping them onto the networks using top 100,000 most confident links, and 1st-order neighbor node gain, expressed as the ratio of non-query nodes that are 1st-order neighbors to the query genes, and the number of query genes. (C) Similarity between the networks expressed as Jaccard index (JI) and Szymkiewicz–Simpson (SS) coefficient for nodes and links. (D) Heat map showing the performance gain values for the 20 top and bottom performing ORPHANET genesets in FunCoup vs HumanNet/STRING. These genesets were selected after sorting by Performance Gain fold-change, which is computed as the difference in performance gain in FunCoup 6 and the maximum performance gain with STRING or HumanNet. The gene set names were shortened for visualization purposes. (E) The FunCoup 6 subnetwork of Usher syndrome type 1 (i.e. USH1K, CDH23, USH1C, USH1G, MYO7A, PCDH15, USH1E, USH1H, CIB2, ESPN) with overall highest performance gain in FunCoup 6. The subnetwork is composed of 22 genes and 41 links when querying the full network (PPV ≥ 0.85) with genes as a group, with expansion step of 1 and 15 nodes per expansion step. USH1E, USH1H, and USH1K are not in the network. (F) The FunCoup 6 subnetwork of Cystic fibrosis (i.e. SERPINA1, SLC26A9, SLC6A14, SLC9A3, CEACAM3, CEACAM6, EDNRA, GSTM3, HMOX1, GCLC, HFE, CFTR, TGFB1, DCTN4, CLCA4, STX1A, KCNN4, SLC11A1, MIF), composed of 34 genes and 89 links. The size of the nodes is proportional to the degree of connectivity. The thickness of the links is proportional to the confidence.

To investigate the reasons behind the different performances in our benchmark, we analyzed network properties of the ORPHANET gene sets. We restricted the following analysis to the benchmark using only top 100,000 links, given that most networks have best average performances at this cutoff. FunCoup 5 has the worst performance, and that is because of a significantly lower genome coverage at high confidence links. Gene sets with low performance gain in an RWR test tend to be those that are fairly well connected to the rest of the network. This means that these gene sets experience a significant increase in nodes when adding first-order neighbors to the ORPHANET subnetwork. Specifically, if we measure the gain in terms of nodes when adding first neighbors to the query set, measured as ratios of non-query nodes and query nodes, we observe that FunCoup 6 has the lowest median gain of nodes at all cutoffs (Figure 4B and Supplementary Figure 9). This suggests smaller expansion and more specific connections, which benefits the RWR by preventing the flow from dispersing too widely across the network. See Supplementary Figure 10 for an analysis of other ORPHANET subnetwork properties.

We also measured how similar the networks are in terms of common nodes and links (Supplementary Figure 11). HumanNet v3 (1,125,484 links) and STRING v12 (6,857,369 links) are the most similar networks, with STRING v12 including about 60% (Szymkiewicz–Simpson (SS) coefficient of 0.58) of the links present in HumanNet v3. FunCoup 6 includes about half of the links in FunCoup 5 (SS=0.48) and 40% of the links present in HumanNet v3. If we compare the networks using only top 100,000 links, overall FunCoup 6 has the most different connections, with a JI of 0.08, 0.13 and 0.14 with HumanNet v3, STRING v12 and FunCoup 5 (Figure 4C).

To highlight the fact that networks differ and are therefore complementary, with gene sets performing differently across them, we isolated the subnetworks with the highest and lowest average performance gain for FunCoup 6 versus the maximum performance gain for either STRING v12 or HumanNet v3. Figure 4D shows the performance gain values for these top and bottom 20 ORPHANET genesets, illustrating the variability in performance across different networks.

To visually distinguish between a well-performing subnetwork and a poorly performing one, we compared the in FunCoup 6 best-performing gene set (Usher syndrome type 1) with the worst-performing gene set (Cystic fibrosis) on average (Figure 4E and 6F). The difference in performance gain between the two gene sets can be attributed to network structure and the distribution of gene importance. In Usher syndrome type 1 the query genes are central and highly connected, facilitating effective network propagation. This structure allows the network to efficiently prioritize the relevant genes. Conversely, in Cystic fibrosis the genes are highly connected but primarily to other genes with even higher degrees. Additionally, some nodes in this gene set are quite disconnected from the overall network, further diminishing the effectiveness of network propagation and resulting in lower performance.

### Orthology-transferred networks for 618 species

In FunCoup 6, we enabled network queries for all species in InParanoiDB 9 by transferring the network from the closest of the primary FunCoup species using InParanoiDB 9 orthologs. To evaluate the orthology-transferred networks, we compared six species, grouped into three pairs of closest species: *B. subtilis - E. coli, S. cerevisiae - S. pombe, and M. jannaschii - S. solfataricus*. The networks of these species were compared with those generated by the evidence-based approach (Supplementary Figure 12), revealing Jaccard Indices up to 0.7 and overlap coefficients up to 1.0 of nodes and links, indicating a close approximation despite the lower coverage of the transferred networks. An exception was observed for *S. pombe*, which had higher link coverage when transferred from *S. cerevisiae* orthologs, likely due to the large amount of data available for the latter.

### The website

The latest update to FunCoup 6 introduces several key improvements to enhance user experience and functionality. The website frontpage has been completely redesigned for a more intuitive user experience, featuring a condensed and more intuitive advanced search function. To address ambiguities in the gene names in the query set, we first perform a database search that identifies any genes that could not be found or have ambiguous names. An intermediate page then lists these genes, allowing users to select the intended genes. The network view utilizes the D3.js library, now fully integrating with the interaction table, so hovering over table rows highlights corresponding links in the network. Additionally, the side panel next to the network enables extensive customization options, allowing users to modify node and link colors, apply filters or colors to pathways and tissues when available, and adjust the query ID, edge widths, and node sizes. Furthermore, a new tab has been added to visualize KEGG pathway enrichment analysis performed using ANUBIX and EASE. These methods were selected because ANUBIX and EASE have demonstrated top speed and performances in our previous benchmark (Buzzao et al. 2024).

## DISCUSSION AND CONCLUSIONS

We present the sixth release of the FunCoup database of functional association networks, where we introduce several methodological advancements and tools designed to enhance network inference and analysis. Key improvements in FunCoup 6 include significantly expanded gene regulatory link coverage, the development of a novel bin-free naïve Bayesian training framework, and a comprehensive redesign of the website.

The significant increase in the number of links in FunCoup 6 is driven by multiple factors. First, the inclusion of more and new datasets expanded the underlying data, allowing for greater coverage. Second, the novel bin-free Bayesian training framework improved the granularity and accuracy of link predictions. Additionally, the new link confidence approach provides a key advantage by balancing the support of each gold standard across the network, leading to a more homogeneous and varied representation of functional associations. This combination of factors has not only increased the number of links but also improved the biological relevance of the networks, particularly in species like human, where the number of directed regulatory links has increased substantially.

Despite the use of standardized pipelines and normalized enrichment scores, a limitation of the GRG approach for inferring regulatory links is the inherent complexity in comparing peaks from independent experiments. A peak with a score of 1 in one experiment may not be as biologically significant as a similarly scored peak in another one due to differences in experimental conditions, sequencing depth and other technical factors. However, in a Bayesian integration framework like FunCoup, protein pairs are unlikely to form high-confidence links without support from additional experimental evidence, thereby reducing the risk of directed false positive interactions.

We opted for a bin-free naïve Bayesian training framework for network inference due to its balanced advantages in computational efficiency, interpretability, and its capability to integrate diverse omics data. This bin-free implementation enhances resolution, accuracy, and training stability by avoiding the arbitrary discretization of continuous data, allowing for a more precise representation of the underlying biological signals. While advanced methods such as neural networks excel at capturing complex, non-linear relationships, they require significant computational resources and often lack interpretability. In contrast, the naïve Bayesian network inference method, which assumes conditional independence between data sources, provides an effective means of integrating multi-modal omics data. This makes it an optimal choice for handling the diverse and complex nature of biological datasets encountered in FunCoup 6.

To ensure fair comparison in our benchmark, we applied network propagation on interactomes with a fixed number of links, performing a standardized number of random link swaps to introduce controlled variability. This method preserves the original degree distribution and topological properties of the networks, ensuring that observed performance differences reflect true variations in network structure rather than artifacts from the shuffling process. The use of ORPHANET gene sets, which are not biased towards common diseases, further reinforces the validity of the used benchmark. By using ORPHANET instead of DisGeNET or GWAS data as done in a previous benchmark (Wright et al. 2024) our results came out differently, although in that study FunCoup, HumanNet, and STRING were all top-performing networks with relatively similar performance.

In order to improve user experience and functionality, we developed a new website featuring intuitive advanced search, improved query disambiguation, a D3.js-powered network view with customization options, and a new tab for KEGG pathway enrichment analysis using top-performing methods (ANUBIX and EASE). We also upgraded the API, which now offers functionality that mirrors the capabilities of the website, allowing users to perform searches programmatically and receive results in JSON format. Detailed instructions for using the updated API are available on the API page of the FunCoup website.

In addition to the MaxLink tool, the incorporation of the TOPAS algorithm facilitates the identification of disease and drug target modules. Another unique feature of FunCoup is the comparative interactomics tool, which provides insights into network conservation and allows for ortholog alignment or independent species network queries. The bin-free training framework has been applied to 23 primary species, with networks for an additional 618 species in InParanoiDB 9 generated using a novel orthology-based method. Together these enhancements in FunCoup 6 reflect our ongoing commitment to providing a robust and versatile resource for the analysis of functional association networks, facilitating a deeper understanding of complex biological processes and their implications in health and disease.

## DATA AVAILABILITY

All data is available at the FunCoup 6 website.

## ACKNOWLEDGEMENTS

This work was supported by the Swedish Research Council [2019-04095]. Open access funding is provided by Stockholm University. We thank Matteo Canegallo for contributing ideas and preliminary studies that inspired the new bin-free network training; Lukas Steininger for contributing ideas to the website implementation; Anastasiia Mironova for the design of the landing page of the website; Arianna Cazzola for designing the FunCoup logo.

## Conflict of interest statement

None declared.

## Notes

https://funcoup6.scilifelab.se

### Competing Interest Statement

The authors have declared no competing interest.

