## Supplementary Materials for "FunCoup 6: advancing functional association networks across species with directed links and improved user experience"

#### MATERIALS AND METHODS

##### Proteomes

In FunCoup 6, all proteomes from FunCoup 5 were updated using UniProt reference proteomes 2022-02. We added the new species *Mycobacterium tuberculosis*. Identifier mappings necessary for integrating different types were extracted from UniProt ID mapping files.

##### Gold standards

In FunCoup, gold standards serve as proxies for true interactions to assign confidence scores to protein associations, regardless of the evidence type. For this update, we used six gold standards, building on five from previous versions and adding a regulatory category to capture the interactions between transcription factors and target genes. The other gold standards include metabolic and signaling pathways, protein-protein interactions (PPIs), protein complexes, and shared operons. When multiple sources contribute to a gold standard, their union is used. See Supplementary Figure 1 for a comparison of the number of links per gold standard between FunCoup 5 and FunCoup 6.

###### Complex

Protein members of the same complex, according to iRefIndex (v2022-08) (Razick, Magklaras, and Donaldson 2008), Corum (v2022-11) ((Giurgiu et al. 2019), and ComplexPortal (v2023-06) (Meldal et al. 2022a)(Meldal et al. 2022b), were considered functionally related and were included in the complex gold standard. Some complexes included thousands of proteins, likely due to false associations. To minimize the impact of such noise during network training, we limited the size of complexes in the gold standard to a maximum of 100 proteins, following the largest complex size found in Corum.

###### Metabolic and Signaling

Metabolic and signaling links were extracted as fully connected networks between proteins belonging to KEGG (v101.0) metabolic and signaling pathways (Kanehisa and Goto 2000).

###### Operon

Shared operon links were extracted from OperonDB (v2022-02) (Pertea et al. 2009).

###### PPI

Protein-protein interactions (PPI) were extracted from iRefIndex (v2022-08). PPI must be supported by at least two experiments or be present in another gold standard. Similarly to the Complex gold standard, we restricted the focus to physical interactions with no more than 100 interactions per experiment.

###### Regulatory

We introduced the Regulatory gold standard TRRUST (v2) (Han et al. 2018) RegNetwork (v2019-04) (Liu et al. 2015), RegulonDB (v2022-12) (Salgado et al. 2024) and Yeastract (v2022) (Teixeira et al. 2018) to capture transcription factor regulatory interactions for (Liu et al. 2015). We used the union of TRRUSTv2 and RegNetwork for human and mouse links.

##### Evidences

FunCoup 6 has ten independent evidences, to capture functional association at multiple omics levels. Data is collected from public online databases and resources. For each data type, a scoring metric is used to measure the strength of association. Raw scores are subjected to transformation and normalization before entering the database.

**DOM**

DOMain-domain interaction (DOM) is based on pre-computed scores from UniDomInt (v1.0). The UniDomInt score indicates support across source databases. Domain interactions were mapped to protein pairs using Pfam (v35.0). DOM is computed as the ratio of the number of scored domain pairs $m$ to the total possible pairs $N$, multiplied by the average of scores, weighted by the log of the sum of the degrees of the two interacting domains (Eq. 1). $N$ is calculated as the product of the number of Pfam domains in $A$ and $B$.

$$DOM(A, B) = \frac{m}{N} \times \frac{\sum_{i=1}^{m} \frac{1}{log(d_a + d_b)} UniDomInt(a,b)}{m} \tag{1}$$

**GIN**

Genetic INteraction (GIN) was measured as a Spearman correlation of genetic interaction profiles as reported in BioGRID (v4.4.219), with minimum 5 valid pairs. This cutoff was chosen to avoid spurious correlations often caused by low-coverage data, which can lead to multimodal, discrete distributions. A genetic interaction profile represents how a gene interacts with many other genes. For example, if Gene A is mutated and tested with Genes B, C, and D, the results (e.g., how much the double mutations affect growth) form Gene A's profile. If Gene B has a similar profile (similar effects with Genes C, D, etc.), it suggests that Genes A and B might be functionally related.

**GRG**

Gene ReGulation (GRG) was extracted from pre-computed Browser Extensible Data (BED) narrowPeak files in ENCODE (v131.0) for human, mouse, worm and fruit fly. We used the enrichment score of Irreproducibility Discovery Rate (IDR) thresholded peaks that have undergone standardised ENCODE (human, mouse) or Robert-Waterston (mouse, fly) processing pipeline (Hitz et al. 2023). To accommodate for technical variability, each dataset was normalized and then intersected with others. Redundant interactions were sorted by maximisation of enrichment score. Pybedtools (v0.9.1) was used to handle gene annotations of detected peaks. Genomes GRCh38 and mm10 were downloaded from ENCODE, ce11 and dm6 were downloaded from Ensembl.

**MEX**

mRNA-coEXpression (MEX) was computed as a Spearman correlation of expression profiles for healthy samples (with minimum 3 valid pairs). Data was extracted from GEO and EBI Expression Atlas. Only pairs with absolute correlation larger than 0.5 were retained, and then min-max scaled between 0 and 1. MEX was not computed for homolog pairs. In FunCoup 6 we used DIAMOND and considered homologs those with default E-value below 0.001. We made a comparison of the number of homologs found with Diamond at bit score above 100 (*i.e.* which is similar to Blast bit score) and E-value below 0.001 (Supplementary Figure 2).

**MIR**

microRNA Regulation (MIR) was computed as the Jaccard Index of microRNA regulating each protein pair, retaining only positive values. Here, we used previous data (Betel et al. 2008).

**PEX**

Protein co-EXpression (PEX) was measured on PaxDB (v5.0) quantitative mass spectrometry (QMS) data in two ways, by measuring the correlation profile over tissues (with minimum 5 valid pairs), and by measuring the Jaccard index of 25% top tissues each protein pair is expressed in, retaining only positive values, as it was previously done in FunCoup with the evidence QMS.

**PHP**

Phylogenetic Profile Similarity (PHP) was assessed using phylogenetic profile data from all 640 species of InParanoiDB 9. An orthophylogram and a distance matrix were constructed using orthologs from the InParanoid species, where the distance between two species was computed as one minus the average fraction of orthologous proteins in both directions, i.e. 1 - [(number of orthologous proteins in species A / total number of proteins in species A) + (number of orthologous proteins in species B / total number of proteins in species B)] / 2. Subsequently, a profile was generated for each protein in the FunCoup species, with the presence of an ortholog represented by 1 and the absence by 0. To determine PHP scores for a specific species, the distance matrix was employed to construct a neighbour-joining tree rooted at the species of interest. For each species, and for every pair of proteins, we used the phylogenetic profiles to calculate the PHP score. This score was derived from the logarithm of the ratio between a "positive score" and a "negative score". The positive score represents the total branch length of the subtree encompassing all species that have orthologs for both proteins, divided by the total branch length of the entire tree. Conversely, the negative score represents the total branch length of the subtree containing species where only one of the proteins has an ortholog, divided by the total branch length of the entire tree. We use branch length to weight the conservation of gene pairs through evolution, with the idea that deeper, more ancient conservation represents a stronger signal of functional association than more recent conservation.

**PIN**

Physical INteraction (PIN) was computed as the weighted average of the number of publications *n* supporting the interaction, as listed in IrefIndex (v2022-08), with weights based on the logarithm of the interactions reported in each publication $PMID_i$, down-weighted by the logarithm of the total number of connections of proteins *A* and *B* (Eq. 2). The degree $d_A$ represents the total number of interactions that protein *A* has in IrefIndex ($d_B$ for protein *B*), and these are used to increase the specificity of the detected interactions by accounting for proteins that have a high number of interactions.

$$PIN_{A,B} = \frac{\sum_{i=1}^{n} 1/log(1+|PMID_i|)}{n} \times \frac{1}{log(d_A + d_B)} \quad (2)$$

**SCL**

SubCellular Localization (SCL) was measured as semantic similarity of Gene Ontology (GO) keywords (v2023-03). To compute similarity scores we used the Wang et al. graph-based method (Wang et al. 2007), which uses the topology of the GO Directed Acyclic Graph structures.

**TFB**

Transcription Factor Binding profile (TFB) was measured as the Jaccard index of transcription factors binding to each protein pair, retaining only positive values. Data was downloaded from TFLink (v1.0) (Liska et al. 2022).

#### Orthology

Evidence types GIN, GRG, MEX, MIR, PEX, PIN and TFB are eligible for evidence transfer using orthologs, meaning that the species specific evidence of those types are transferred using orthologs from InParanoiDB 9 (Persson and Sonnhammer 2023) to each of the FunCoup species, and is used as an additional source of evidence. Scores for many-to-one relationships (e.g., A1, B1, and C1 in species 1 are all orthologs of A2 in species 2 and pair with D1, an ortholog of D2) were averaged into a single score to avoid duplicates. Evidence types DOM, PHP, and SCL are not transferred between species because they already contain cross-species information.

In FunCoup 6 we introduced the possibility to query networks for all 640 species available in InParanoiDB 9. This is done by transferring entire networks for each InParanoid species from their closest species among the 22 FunCoup networks using InParanoiDB 9 orthologs. The closest FunCoup species for each of the InParanoid species was obtained by taking NCBI taxonomy lineage information, and finding the FunCoup species closest to it in the taxonomy tree. For species where taxonomy information was missing from NCBI, or where ties occurred, i.e. a species was equally close to multiple FunCoup species, the species with the shortest distance in the orthophylogram for InParanoiDB9 was used. We used the unweighted pair group method (UPGMA) to build a tree from distances calculated as 1 minus the fraction of orthologous proteins averaged over both directions for all InParanoiDB 9 species pairs. When transferring a network, each link in the original network was transferred to all possible pairs of orthologs in the target species, keeping the original link score. All transferred networks are available for download from the FunCoup website, and the transferred species can be queried in the FunCoup website using UniProt IDs. All query types and options are available when querying for a transferred species, but as KEGG pathway information is not available for all transferred species, pathway enrichment was deactivated. The orthology-based inference approach was evaluated by comparing the networks of the species *B. subtilis, E. coli, M. jannaschii, S. cerevisiae, S. pombe,* and *S. solfataricus*, with the evidence-based approach. For each of the six target species, an independent network was trained by deactivating orthology transfer from the target species evidence, and then an orthology-transferred network was inferred from that. The network comparisons were made in terms of Jaccard Index (JI) and overlap coefficient (OC) similarity of nodes and links between the evidence-based and orthology-transferred networks.

#### Bin-free network training

FunCoup 6 uses a Bayesian framework to extract likelihoods of functional associations from high-throughput data and gold standards. This involves deriving odds of likelihoods through link randomization. Positive links within the database, from the species under training and all other available species, are then filtered for each gold standard. Likelihoods are extracted for each evidence-gold standard combination, using kernel density estimation with Gaussian kernel and the Silverman's Rule (Silverman 2018) for bandwidth optimization. The likelihood ratio is computed by calculating the ratio of positive and negative likelihoods over a common range of raw scores. Polynomial regression models the likelihood ratio distribution within a constrained range, typically up to the 98th percentile, excluding outliers. Successful training requires a dataset with a range of $10^3$-$10^6$ observations, and the polynomial regression must meet specific criteria, including an $R^2$ value exceeding 0.9 and a polynomial degree between 2 and 4. For each gold standard network, a Final Bayesian Score (FBS) is extracted as a sum of the log likelihood ratios between all evidences that successfully passed the training requirements. The new confidence estimation method involves assessing Positive Predictive Value (PPV) across increasing Final Bayesian Scores (FBS). In this method, positive examples consist of gold standard links with positive FBS, and negative examples are sampled among the rest, for 30 times, to be as many as the positive examples. The PPV was computed as TP / (TP+FP) over a range of 1000 threshold values using the precision_score function from the python sklearn.metrics library. In such a way, an FBS of 0 would correspond to a PPV of 0.5. The PPV confidence for all other protein pairs is then calculated by fitting a logistic function to the PPV vs FBS relation, providing an empirical measure of the likelihood of functional associations within the network. For each species, the final FunCoup network corresponds to the union of the gold standard networks with maximum PPV. Only links with PPV≥0.85 are kept. In the updated scoring system of FunCoup, we introduce the concept of gold standard-based PPV (gsPPV) to integrate gold standard links more effectively. gsPPV values were assigned based on the number of gold standards supporting a link, with the values ranging between the minimum and maximum PPV stored in the database. Specifically, gsPPV was set to 0.85 for links supported by one gold standard, 0.90 for two, 0.95 for three, and 1 for more than three. This is because most species have support from up to four gold standards (Complex, Metabolic, PPI and Signaling). However, if a link has a PPV value, estimated based on evidence, that is exceeding the gsPPV value, the gsPPV value is discarded.

#### New redundancy schema

An additive sequential redundancy schema was implemented to reduce the redundancy effect of MEX on the FBS. For each pair *A,B* LLRs are ranked in decreasing order and summed up, down-weighted by a decreasing factor *w,* so that $w_i = w_i * w_{i-1}$. *w* is computed as follows: $w_{e,k} = \alpha[1 - max(0, r_{e,k})]$, with $\alpha = 0.7$ as it was optimized for previous versions of FunCoup. In this case, $r_{e,k}$ represents the Spearman correlation coefficient between dataset $e$ and $k$ within the evidence MEX and it was used to estimate the degree of redundancy between them. The correction parameter $\alpha$ accounts for noise in the data.

#### Network Benchmark

To evaluate the performance of FunCoup 6, we compared its human network against FunCoup 5, HumanNet v3, and STRING v12 using ORPHANET gene sets that contain at least 10 genes. To ensure compatibility with each network's gene vocabulary, and convert gene and protein identifiers to Gene Names, we utilized the UniProtKB UP000005640 file, v2021-02 for FunCoup 5 and v2022-02 for FunCoup 6, and HumanNet v3. For STRING v12, we used 9606.protein.aliases.v12.0.txt. For each of the 110 resulting distinct gene sets, we split disease-associated genes into two halves, mapping them to networks and evaluating their recovery using Random Walk with Restart (RWR). The RWR algorithm was executed using the dnet::dRWR function from the R (version 4.3.2) package dnet, with starting probability of 0.75 from half of the genes in each geneset. Performance was assessed by computing the Area Under the Receiver Operating Characteristic (AUROC) curve. True positives were defined as the remaining half of the disease-associated genes for the investigated disease, while all remaining genes were classified as false positives. To avoid bias in the results, we created 30 distinct splits of each of the 110 gene sets. We quantified the Performance Gain $PG$ to evaluate how the real network compares to randomized networks. The benchmark was conducted under five distinct scenarios, by selecting the top 100, 250, 500, 750 thousands and 1 million of the highest confidence links for each network. Randomization of networks was achieved using the igraph::rewire() function with keeping_degseq() mode. This randomization process preserves the node degree distribution of the networks and was performed without allowing for self-interactions (loops=False) and with the number of edge swaps equal to the size of the network.

$$PG(X) = \frac{AUROC_{S_{30}}(X) - AUROC_{RS_{900}}(X)}{AUROC_{RS_{900}}(X)} \qquad (3)$$

Where $X$ is one gene set in the ORPHANET library; $PG$ is the performance gain and $AUROC$ is the Area Under the Receiver Operating Characteristic curve; $AUROC_{S_{30}}(X)$ is the median AUROC computed for 30 splits of the geneset $X$; $AUROC_{RS_{900}}(X)$ is the median AUROC computed for 30 splits of the geneset $X$, under 30 independent randomizations of the network.

#### The website

#### Implementation of FunCoup 6

FunCoup 6 was built in Python (v3.10.8) in a version-controlled conda (v23.9.0) environment, using the Django (v4.1.1) framework. All data is stored in a PostgreSQL (https://www.postgresql.org/) database. Bootstrap (https://getbootstrap.com/) frontend framework was used as a base for the frontend components, and D3.js (https://d3js.org/) was used as a base for drawing a network for each query input. The FunCoup 6 application is run and deployed in a Docker (https://www.docker.com/) container, managed with Docker Compose (version 2.17.2).

#### Network search algorithms

In FunCoup 6, the search capabilities have been significantly expanded beyond the previous two methods: querying genes as a group and querying each gene independently. The default search mode is group search. When using this mode, the database is queried with a set of genes, and the top N genes most connected to the query set are added. Activating the option to prioritize common neighbors increases the likelihood of selecting genes that interact with multiple query genes. In the independent search mode, the database is queried separately for each gene, and at each expansion step, the top N most connected genes for each individual query gene are added. Today, users can also use Maxlink and TOPAS for network queries. Maxlink identifies candidate genes linked to query genes, retaining those with significant interactions according to a hypergeometric test. It was previously accessible through a separate search page that used the now-deprecated Maxlink website, and has been fully integrated into the FunCoup framework. This integration allows Maxlink to be used in conjunction with all other search filters, settings, and restrictions available in FunCoup. These include comparative interactomics and querying specific gold standard networks and evidence types. In addition to Maxlink, the TOPAS algorithm has been added as a new network querying option. TOPAS is designed for detecting modules within networks and has proven effective in identifying biologically relevant disease modules. It can be used for queries involving multiple genes and can be combined with other search settings, including comparative interactomics. The comparative interactomics feature has also been redesigned to support different search strategies. Users can now choose to align orthologs only, which identifies the orthologs of the query network. Alternatively, they can independently query each species network and subsequently draw orthology relationships. There is also an option to use species-specific evidence exclusively.

#### Pathway enrichment analysis

Pathway enrichment analysis is a tool for interpreting gene sets that we can use to identify key biological functions and processes impacted by a group of genes, such as those belonging to a FunCoup subnetwork. Two newly integrated algorithms, EASE and ANUBIX are available on our website to identify network crosstalk between subnetwork genes and KEGG pathways. We have integrated the EASE algorithm, which belongs to the OVerlap Analysis (OVA) class of Enrichment Analysis (EA) methods. EASE tests the proportion of subnetwork genes in KEGG pathways against a discrete probability distribution model and extracts one-sided P-values. It uses a conservative modification of Fisher's exact test by subtracting 1 from the overlap count. ANUBIX is a Network Enrichment Analysis (NEA) method. NEA methods evaluate the interconnectivity between a query gene set and a pathway within a functional association network. This evaluation employs either a parametric or a permutation-based approach to determine the statistical significance of the observed interconnectivity. ANUBIX assesses the significance of crosstalk using a beta-binomial distribution, tailored to each pathway and query gene set through a degree-aware resampling process. The analysis with ANUBIX can be further explored by seamless redirection to PathBIX (Castresana-Aguirre, Persson, and Sonnhammer 2021), a dedicated web-based application. For these analyses, we use pre-computed statistics on FunCoup networks with confidence link scores above 0.95.

### SUPPLEMENTARY FIGURES

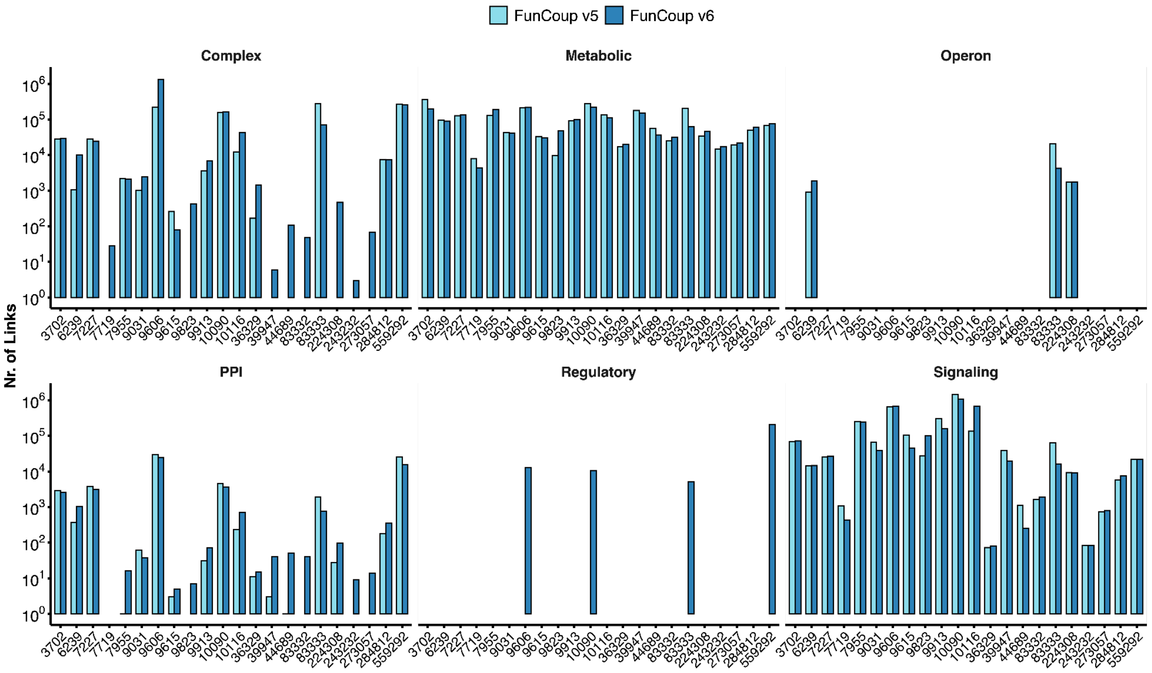


**Supplementary Figure 1 – Number of links per gold standard across FunCoup species, comparing FunCoup 5 and FunCoup 6.** The figure provides a comparison of link counts associated with different gold standards for various species between the two versions.

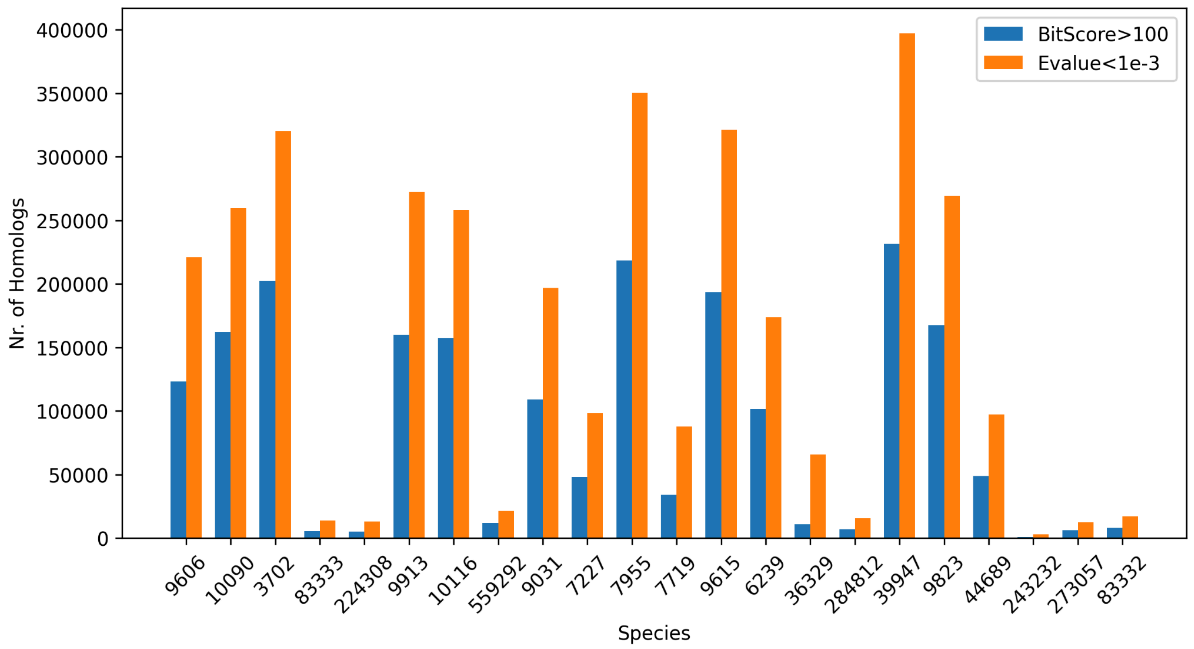


**Supplementary Figure 2 - Comparison of number of homologs between FunCoup versions.** In FunCoup 5 we excluded MEX support for protein pairs that had Blast-based bit scores larger than 100. In FunCoup 6 we consider homologs protein pairs with Diamond-based E-value less than 10e-3. Here we compare the number of homologs between Diamond-based bit score (*i.e.* which is similar to Blast bit score) and E-value.

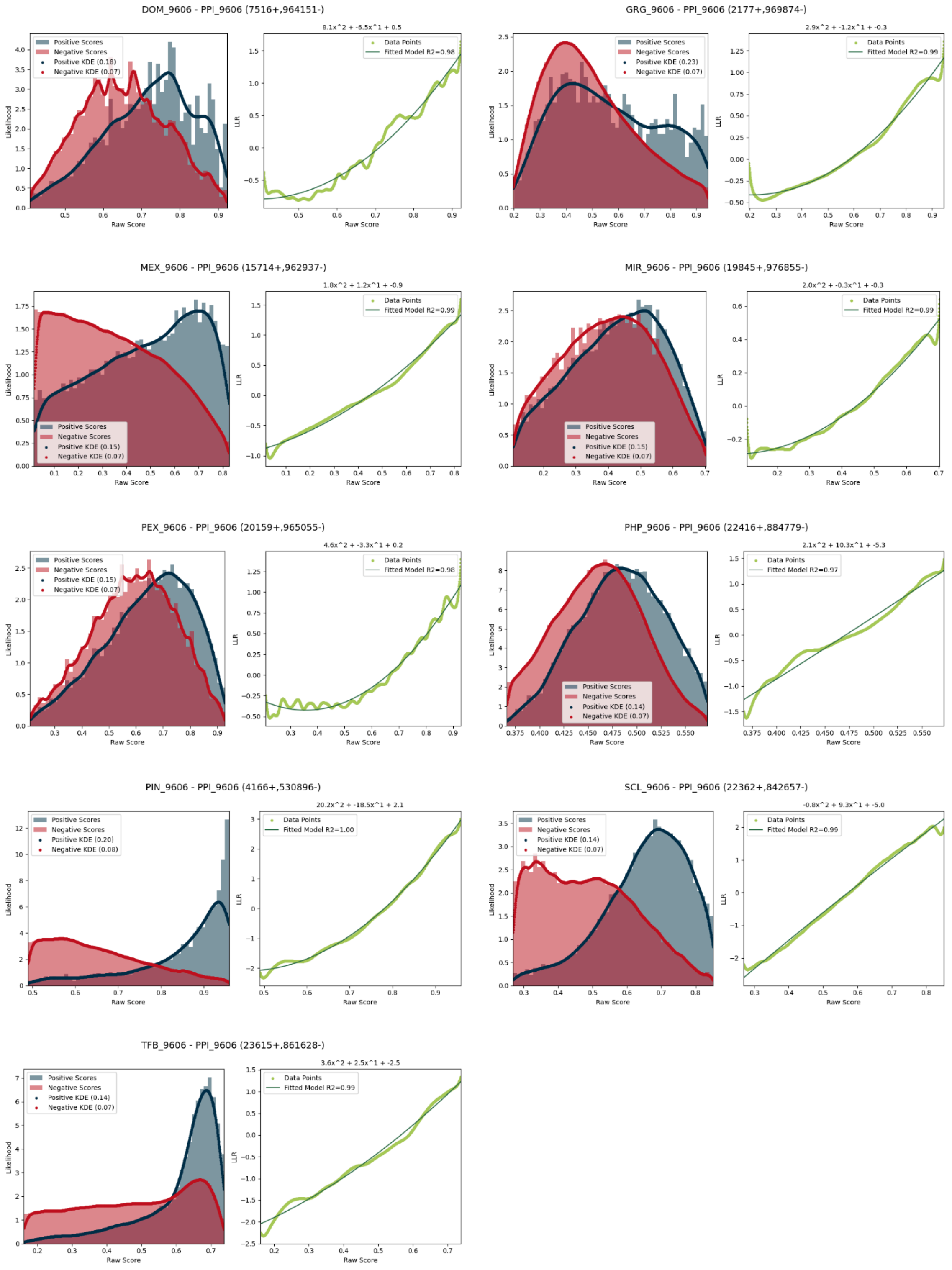


**Supplementary Figure 3 - Inference of log-likelihoods in bin-free naïve Bayesian network training.** Diagnosis plots with Kernel Density Estimation (KDE) of PPI likelihoods (left panel) and polynomial regression of Log-Likelihood Ratios (LLR) (right panel) from 9 out of 10 evidence data in H.sapiens. Training of GIN evidence was unsuccessful with the PPI gold standard due to lack of data (*i.e.* PPI links with GIN score were fewer than 100).

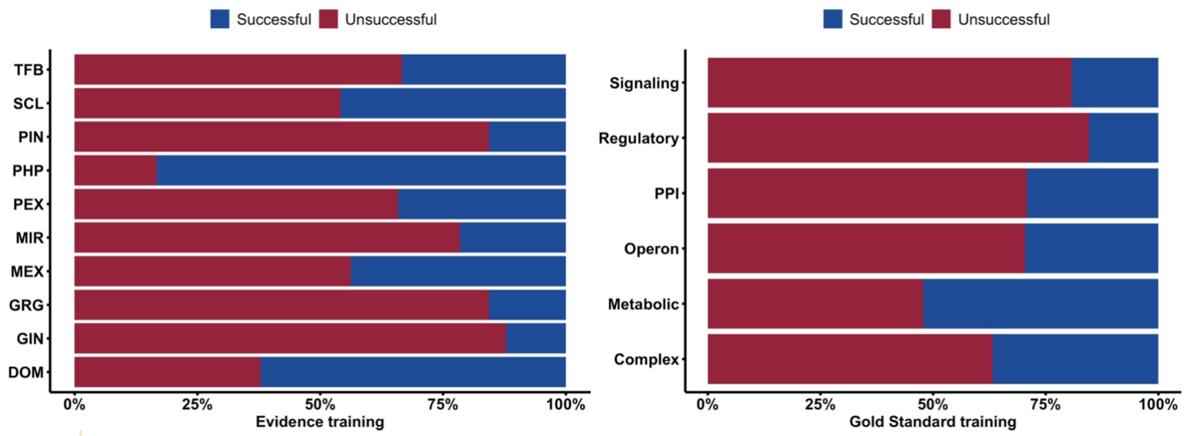


**Supplementary Figure 4 - Success rate of log-likelihood extraction using bin-free naïve Bayesian network training.**

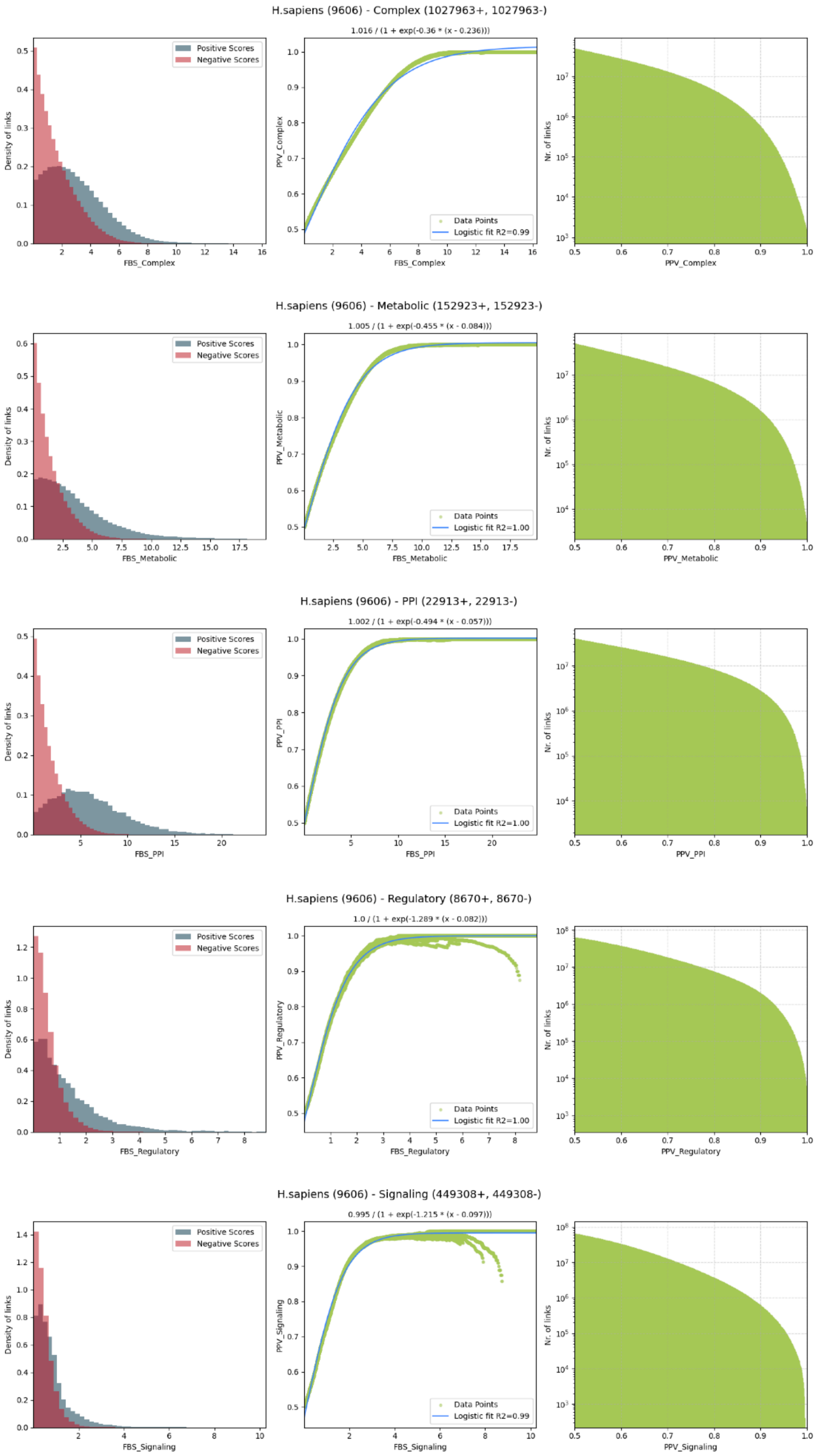


**Supplementary Figure 5 - Inference of PPV confidence.** Diagnosis plots with density of 30x sampled positive and negative examples (left panel), logistic regression of Final Bayesian Score (central panel), and cumulative distribution of number of links (right panel) for all five gold standard networks in H.sapiens.

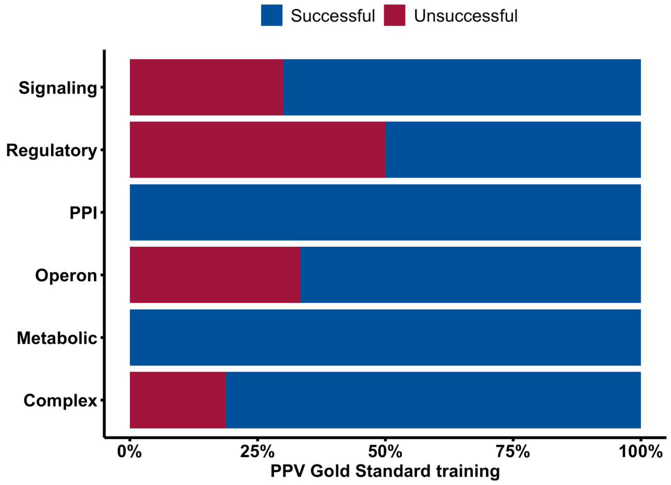


**Supplementary Figure 6 - Success rate of PPV confidence extraction.**

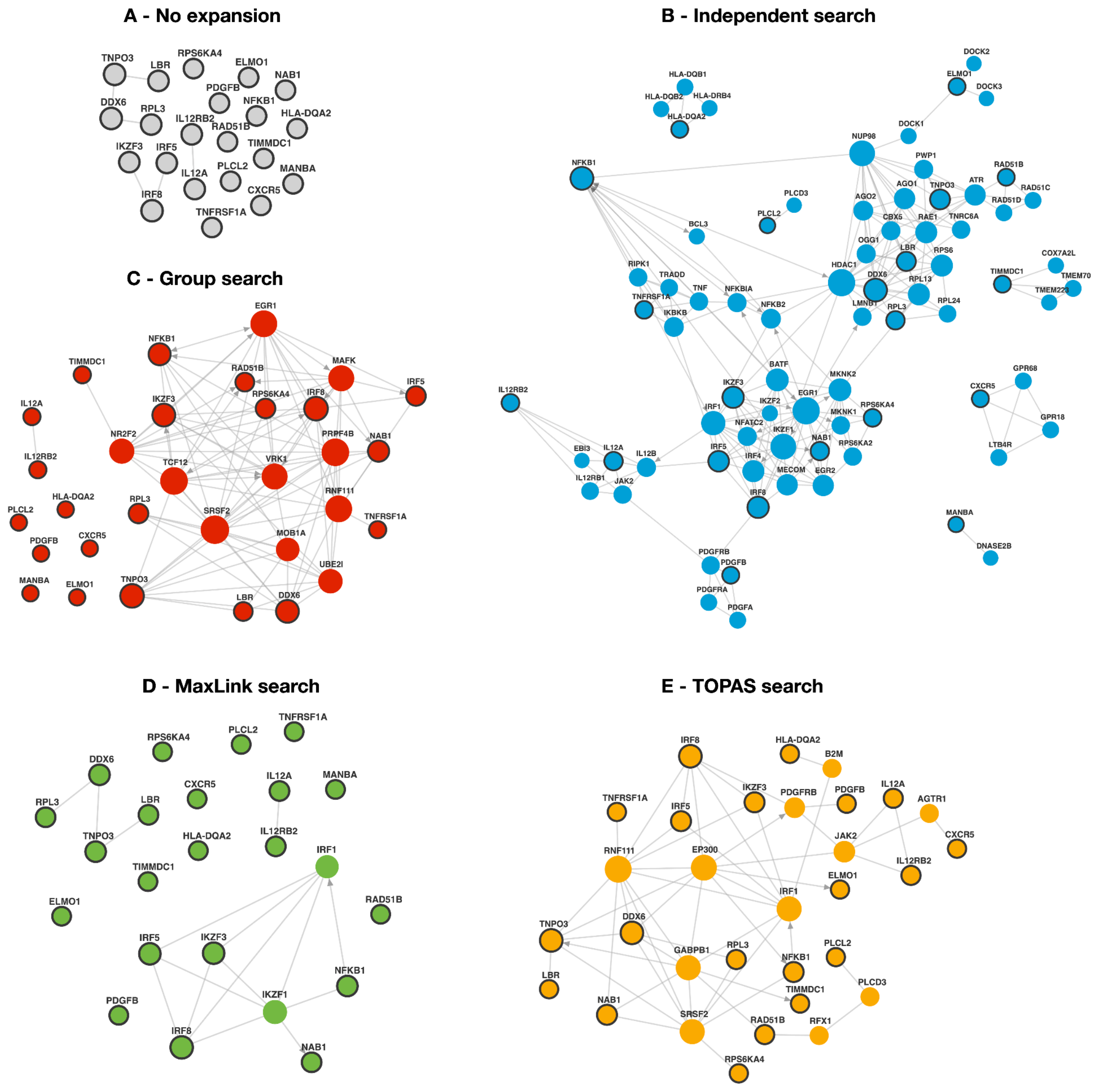

**Supplementary Figure 7 - Subnetworks of *Biliary liver cirrhosis* using four different search algorithms.** The database query (I12R2, NAB1, TIDC1, RA51B, KS6A4, MANBA, PDGFB, DQA2, TNR1A, NFKB1, DDX6, IL12A, CXCR5, RPL3, IRF8, IRF5, LBR, ELMO1, IKZF3, PLCL2, TNPO3) was performed via the website with link and direction confidence thresholds of 0.95 and 1. (A) Only query genes were searched for connections in the network. (B) Genes were queried independently and the top 3 most confident interactions for each individual query gene were added to the network. (C) Genes were searched as a group, with the prioritization of neighbors option activated, and expansion depth of 1 and max 10 neighbors. (D) Gene prioritization via MaxLink, with 15 maximum candidates at a p-value cutoff of 0.05. (E) Disease module detection by TOPAS with 2 maximum allowed connectors in a shortest path between any two query genes. In the network illustration: query genes are highlighted with bold outlines, the size of the nodes is proportional to the degree of connectivity, and nodes are colored using a new website feature.

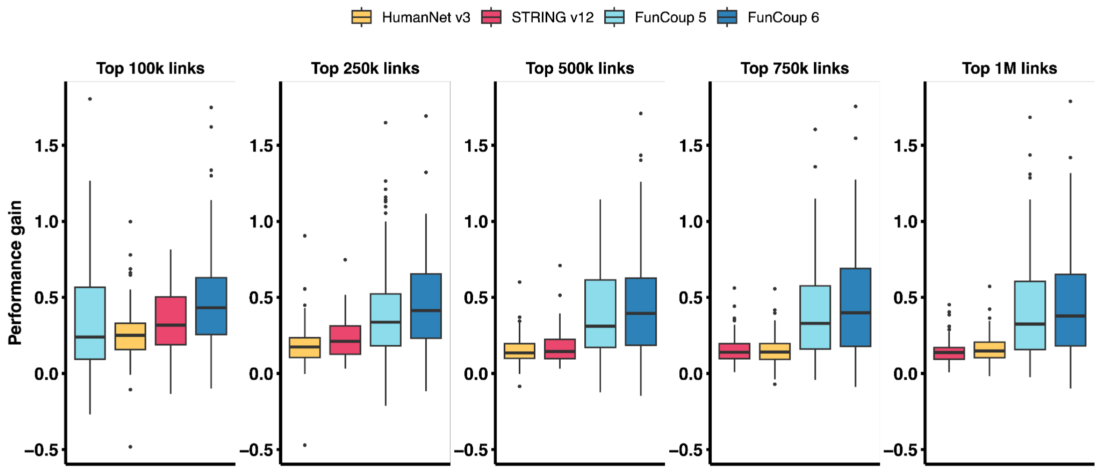


**Supplementary Figure 8 - Performance gain of networks in the ORPHANET benchmark.** Performance gain is quantified as the difference in Area Under the Receiver Operating Characteristic curve (AUROC) between real networks and their null counterparts. The assessment was done using a top fixed number of links: 100,000, 250,000, 500,000, 750,000, and 1.000.000.

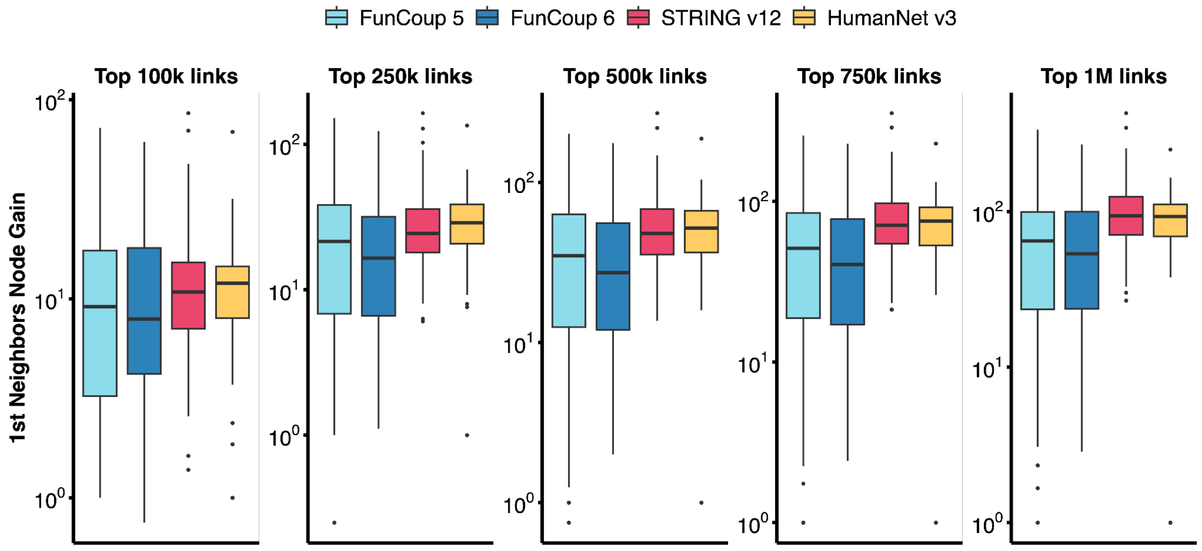

**Supplementary Figure 9 - First-order neighbours nodes gain distribution across benchmarked networks.** The distributions of ORPHANET gene set sizes after mapping them onto the networks using top 100.000, 250.000, 500.000, 750.000 and 1.000.000 most confident links, and 1st-order neighbor node gain, expressed as the ratio of non-query nodes that are 1st-order neighbors to the query genes, and the number of query genes.

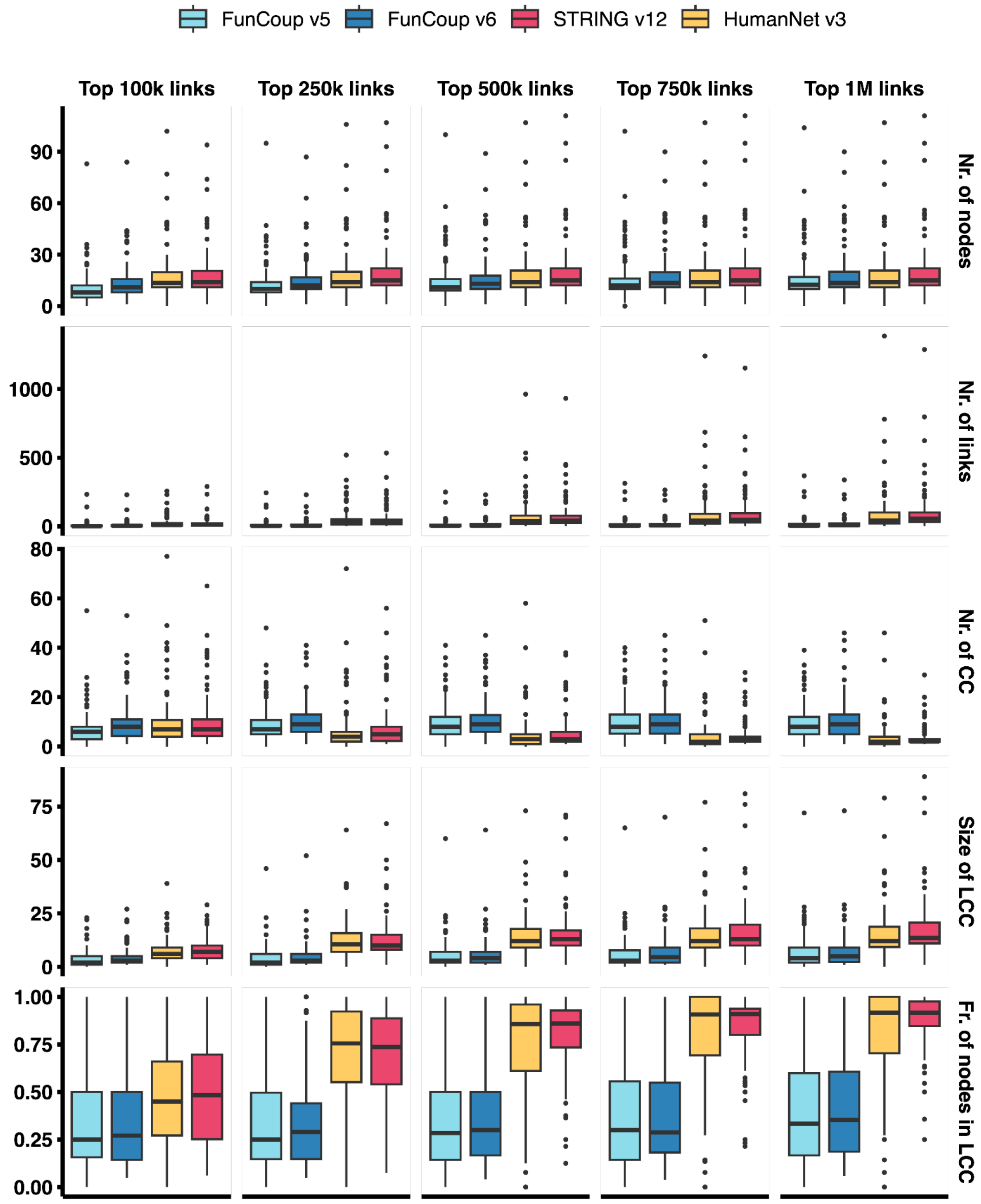


**Supplementary Figure 10 - Overview of ORPHANET gene set properties across all benchmarked networks.** This figure presents key metrics for each network, including the number of nodes, links, connected components (CC), size of the largest connected component (LCC), and the fraction of nodes within the LCC.

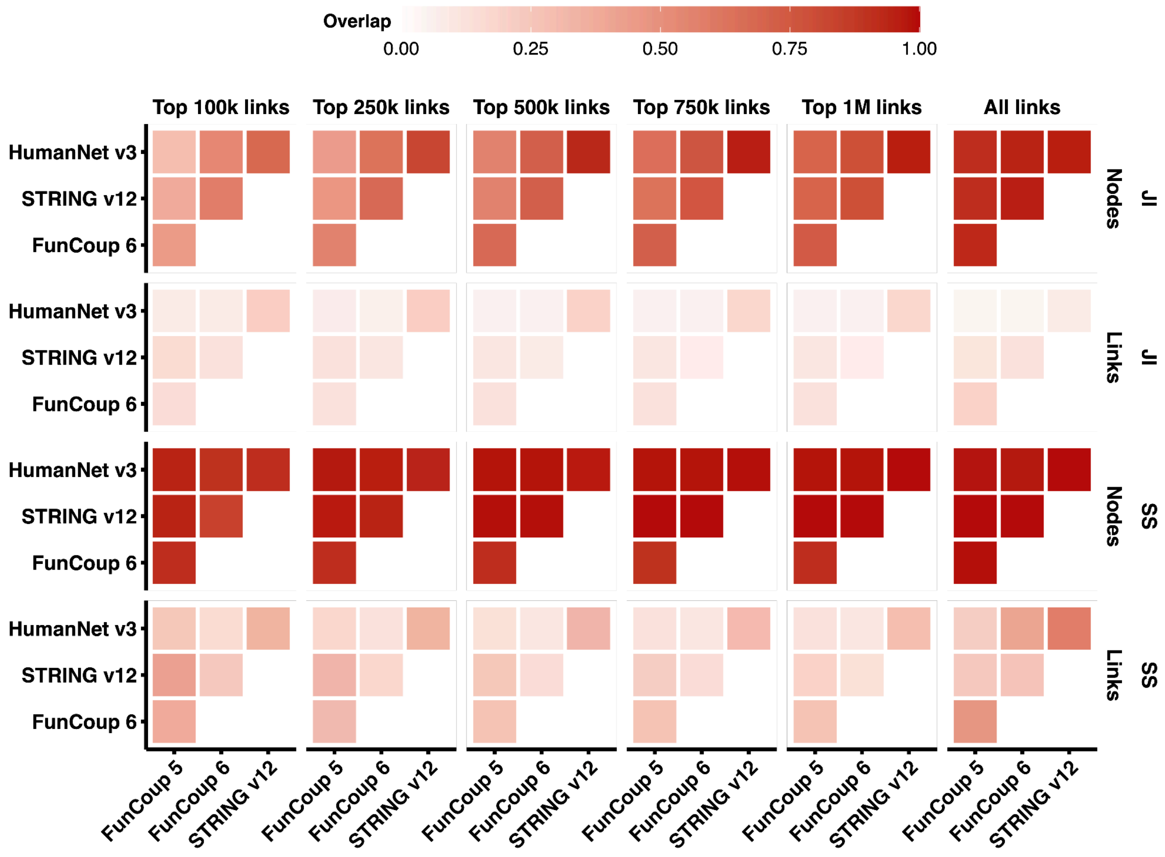


**Supplementary Figure 11 - Benchmarked network similarity.** Similarity between the networks expressed as Jaccard index (JI) and Szymkiewicz–Simpson (SS) coefficient for nodes and links.

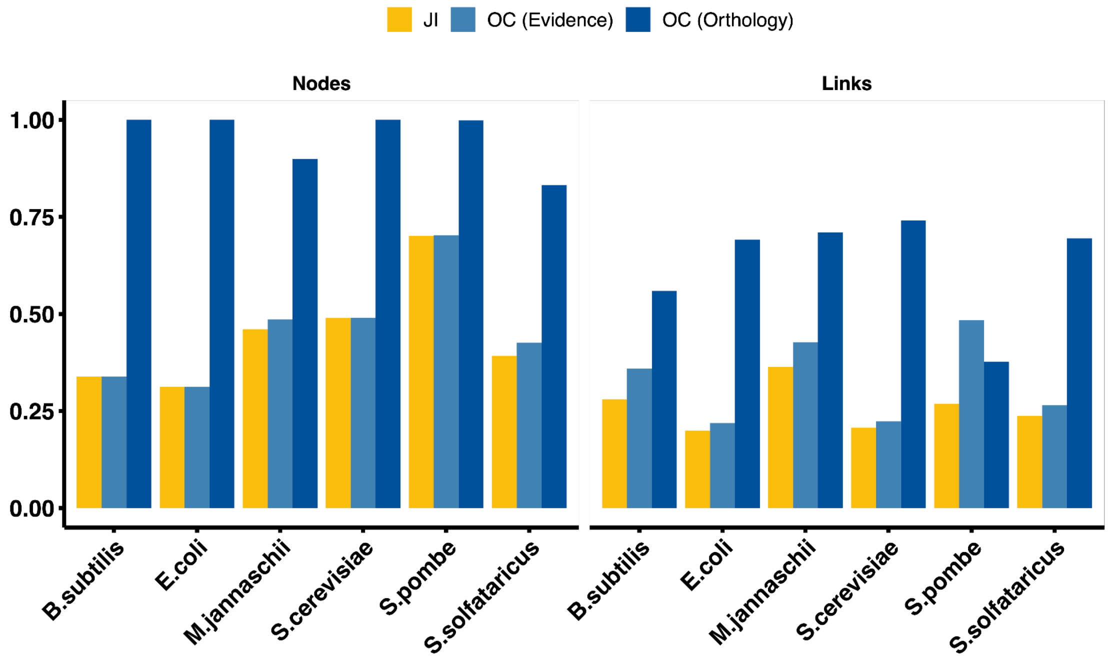


**Supplementary Figure 12. Comparison between evidence-based and orthology-transferred networks.** To assess the quality of the orthology-transferred networks, we compared them against evidence-based networks for six species using the Jaccard Index (JI) and overlap coefficient (OC) for nodes and links. The overlap coefficient is normalized separately for the evidence-based (Evidence) and orthology-transferred (Orthology) networks, resulting in two distinct values

#### SUPPLEMENTARY TABLES

**Supplementary Table 1 - Summary of changes to data source and scoring methods for the FunCoup evidences.** Differences between FunCoup 5 and 6 for all evidences (i.e. DOM, domain interactions; GIN, genetic interaction profile similarity; GRG, gene regulation; MEX, mRNA co-expression; MIR, co-miRNA regulation by shared miRNA targeting; PEX, protein co-expression; PHP, phylogenetic profile similarity; PIN, protein interaction networks; QMS, quantitative mass spectrometry data; SCL, subcellular colocalization; TFB, shared transcription factor binding). Details about the scoring methods can be found in the Methods section.

| | Data source | | Scoring method | |
|---|---|---|---|---|
| Evidence | FunCoup 5 | FunCoup 6 | FunCoup 5 | FunCoup 6 |
| DOM | UniDomInt v1.0 | UniDomInt v1.0 | UniDomInt score | Norm. average weighted UniDomInt score |
| GIN | Costanzo et al. 2010-2017 | BioGRID v4.4.219 | pre-computed scores | Spearman correlation |
| GRG | — | ENCODE v131.0 | — | Norm. maximum peak Enrichment Score |
| MEX | GEO | EBI Expression Atlas, GEO | Absolute Spearman correlation | Absolute Spearman correlation ≥ 0.5 |
| MIR | microRNA.org v2010, MirTarBase | microRNA.org v2010 | Jaccard-index like | Jaccard-index > 0 |
| PEX | HPA v19 | PaxDb v5.0 | Weighted mutual information | Jaccard-index > 0, and Spearman correlation |
| PHP | InParanoiDB v8.0 | InParanoiDB v9.0 | Ortholog ratio | Ortholog log ratio |
| PIN | iRefIndex v16 | iRefIndex v2022-08 | Average weighted PubMed publications | Norm. average weighted PubMed publications |
| QMS | PaxDb v4.1 | — | Jaccard-index like | — |
| SCL | GeneOntology v2022-03 | GeneOntology v2023-03 | Weighted mutual information | Graph-based semantic similarity |
| TFB | ENCODE, Yeastract 2011-2016 | TFLink v1.0 | Jaccard-index like | Jaccard-index > 0 |

Razick, Sabry, George Magklaras, and Ian M. Donaldson. 2008. “iRefIndex: A Consolidated Protein Interaction Database with Provenance.” BMC Bioinformatics 9 (September):405.

Salgado, Heladia, Socorro Gama-Castro, Paloma Lara, Citlalli Mejia-Almonte, Gabriel Alarcón-Carranza, Andrés G. López-Almazo, Felipe Betancourt-Figueroa, et al. 2024. “RegulonDB v12.0: A Comprehensive Resource of Transcriptional Regulation in E. Coli K-12.” Nucleic Acids Research 52 (D1): D255–64.

Silverman, Bernard W. 2018. Density Estimation for Statistics and Data Analysis. Routledge.

Teixeira, Miguel C., Pedro T. Monteiro, Margarida Palma, Catarina Costa, Cláudia P. Godinho, Pedro Pais, Mafalda Cavalheiro, et al. 2018. “YEASTRACT: An Upgraded Database for the Analysis of Transcription Regulatory Networks in Saccharomyces Cerevisiae.” Nucleic Acids Research 46 (D1): D348–53.

Wang, James Z., Zhidian Du, Rapeeporn Payattakool, Philip S. Yu, and Chin-Fu Chen. 2007. “A New Method to Measure the Semantic Similarity of GO Terms.” Bioinformatics 23 (10): 1274–81.
